# Haemosporidian taxonomic composition, network centrality and partner fidelity between resident and migratory avian hosts

**DOI:** 10.1101/2021.06.02.446816

**Authors:** Daniela de Angeli Dutra, Alan Fecchio, Érika Martins Braga, Robert Poulin

**Author notes:** **Correspondence**: Daniela de Angeli Dutra.

## Abstract

Migration can modify interaction dynamics between parasites and their hosts with migrant hosts able to disperse parasites and impact local community transmission. Thus, studying the relationships among migratory hosts and their parasites is fundamental to elucidate how migration shapes host-parasite interactions. Avian haemosporidian parasites are some of the most prevalent, diverse, and important wildlife parasites, and are also widely used as models in ecological and evolutionary research. Here, we contrast parasite taxonomic composition, network centrality and partner fidelity among resident and non-resident hosts using avian haemosporidians as study model. In order to evaluate parasite taxonomic composition, we performed permutational multivariate analyses of variance to quantify dissimilarity in haemosporidian lineages infecting different host migratory categories. Additionally, we ran multilevel Bayesian models to assess the role of migration in determining centrality and partner fidelity in host-parasite networks of avian hosts and their respective haemosporidian parasites. We observed similar parasite taxonomic composition and partner fidelity among resident and migratory hosts. Conversely, we demonstrate that migratory hosts play a more central role in host-parasite networks than residents. However, when evaluating partially and fully migratory hosts separately, we observed that only partially migratory species presented higher network centrality when compared to resident birds. Therefore, migration does not lead to differences in both parasite taxonomic composition and partner fidelity. However, migratory behavior is positively associated with network centrality, indicating migratory hosts play more important roles in shaping host-parasite interactions and influence local transmission.

## Introduction

Migration, i.e. long distance and periodical roundtrip movement of animals between distinct habitats, can alter interaction dynamics among parasites and their hosts by serving as an escape mechanism from some pathogens but also increasing parasite prevalence and richness of certain other pathogens within migrant host species (Altizer et al. 2011; Satterfield et al. 2015; de Angeli Dutra et al. 2021a; Poulin and de Angeli Dutra 2021). Migratory behavior can also modify the availability of hosts for parasites across regions since migrant individuals do not inhabit the same habitat year-round (Bauer and Hoye 2014). At the same time, migrants can represent an opportunity for parasites to increase their distribution worldwide, as infected migrant individuals transport their pathogens through their routes and stopovers, therefore, providing new opportunities for host switching into new environments and resident species (Altizer et al. 2011; de Angeli Dutra et al. 2021b; Poulin and de Angeli Dutra 2021). Indeed, the presence of migratory individuals can also affect local parasite transmission, altering parasite prevalence and richness within resident host communities (Bauer and Hoye 2014; de Angeli Dutra et al. 2021b; Fecchio et al. 2021). However, despite the fact migration can modulate parasite-host interaction, only a few studies have addressed the implications of host migration for parasite ecology and evolution (Poulin and de Angeli Dutra 2021).

Intrinsic characteristics of host-parasite interactions could be altered by host migratory behavior, including traits such as virulence (i.e. pathogenicity level) or partner fidelity, i.e. the species specificity in pairwise host-parasite associations. In this context, network analysis can be a powerful tool to explore the roles of particular species in host-parasite interactions (Runghen et al. 2021). Previous research suggests antagonistic interactions display lower partner fidelity than mutualistic ones, indicating host-parasite interaction networks are evolutionarily malleable (Fortuna et al. 2020) and hosts traits can drive network descriptors (Campião and Dáttilo 2020). Additionally, infecting migratory individuals may pose a challenge to parasites due to the need to adapt to novel resources and conditions, which could lead to looser fidelity among parasites and their migrant hosts. For example, for malaria parasites infecting migratory birds to be transmitted into their hosts’ new habitats, they must be able to infect and complete their cycle in new vector species under distinct environmental characteristics (Valkiūnas 2005). Hence, the exposure of parasites to abrupt environmental and vector changes may impact the ecological and evolutionary relationship between parasites and their migratory hosts since host migrations represent repeated, predictable, and directional selective pressures (Møller and Szép 2011; Poulin and de Angeli Dutra 2021). Therefore, it is essential to study how host shifts between migratory and resident hosts occurring in sympatry and under different environmental conditions can alter parasite-host dynamics. This is necessary to elucidate how parasite life-history traits evolve under repeated and predictable changes.

Avian haemosporidian parasites, i.e. malaria and malaria-like vector borne protozoan parasites, are some of the most prevalent, diverse and studied wildlife pathogens. These parasites are an excellent ecological and evolutionary model to study host-parasite relationships due to their high prevalence, diversity, cosmopolitan distribution and variable levels of specificity to their hosts (Valkiūnas 2005). This is particularly relevant for South America, which harbors the highest diversity of birds, vectors and haemosporidian parasites worldwide (Remsen et al.; Santiago-Alarcon et al. 2012; Ellis et al. 2019). This region is also characterized by great vector abundance and considerable haemosporidian prevalence (Braga et al. 2011; Santiago-Alarcon et al. 2012). Furthermore, avian community composition seems to impact parasite composition as well, with avian community turnover driving both haemosporidian and ornithophilic mosquito turnover across the Amazon region (De La Torre et al. 2021). All those features together make South America an ideal region to investigate ecological and evolutionary dynamics of avian haemosporidian interaction.

For the above reasons, studying the role of host migratory behavior in shaping parasite taxonomic composition (i.e. the set of distinct parasite lineages infecting a given host species), network centrality (i.e. the position a species occupies in the host-parasite interaction network) and partner fidelity is fundamental to understand the impact of host migration on life-history traits for parasites. Here, we hypothesize that resident species show higher partner fidelity to their parasites due to the greater stability of environmental conditions and vector species they face. Additionally, since migrants harbor higher richness of haemosporidians (de Angeli Dutra et al. 2021a) and because the more unstable environmental conditions and vectors they encounter may favor their infection by generalist parasites, we also expect them to occupy more central positions in host-parasite networks. Moreover, since migrants are exposed to more parasite lineages as they visit regions that harbor different parasite communities, our second hypothesis is that parasite taxonomic composition differs between resident and migratory avian hosts species. In this study, we computed and compared partner fidelity and network centrality levels between haemosporidians and their resident and partially and fully migratory avian hosts using Bayesian multilevel models. Further, using permutational multivariate analyses of variance (PERMANOVA) we evaluated whether resident and migratory hosts harbor similar haemosporidian assemblages.

## Methods

### Dataset

All analyses were performed using a dataset comprising ∼15200 individual birds representing 974 avian species. Avian communities were sampled in 85 different localities across seven different South American biomes - Amazonia, Atlantic Rain Forest, Cerrado, Temperate Grassland, Caatinga, Pantanal and Andean Forest (Fig. 1). The birds were sampled from 2005 to 2018 with a subset of those samples having previously been used in published research (Lacorte et al. 2013; Ferreira et al. 2017; Fecchio et al. 2019a, 2020; Anjos et al. 2021) and the rest consisting of unpublished data. This large dataset was combined with data available from MalAvi (http://130.235.244.92/Malavi/) and represents a total of 2758 sequenced parasites representing 752 distinct lineages, all belonging to one of three genera: *Plasmodium, Haemoproteus* and *Leucocytozoon*. Haemosporidian infection was estimated using PCR protocols described previously (Fallon et al., 2003; Hellgren et al., 2004; Bell et al., 2015). All lineages were identified by sequencing a DNA fragment obtained using PCR protocols described by Hellgren et al. (2004) that amplify a cytochrome b fragment of 478 base pairs. Hosts were classified into three migratory categories: (1) resident; (2) partial migrant and (3) full migrant, according to the Brazilian Committee of Ornithology Records - CRBO 2014, Somenzari et al., 2018 and BirdLife International (https://www.birdlife.org/).

**Fig. 1:**
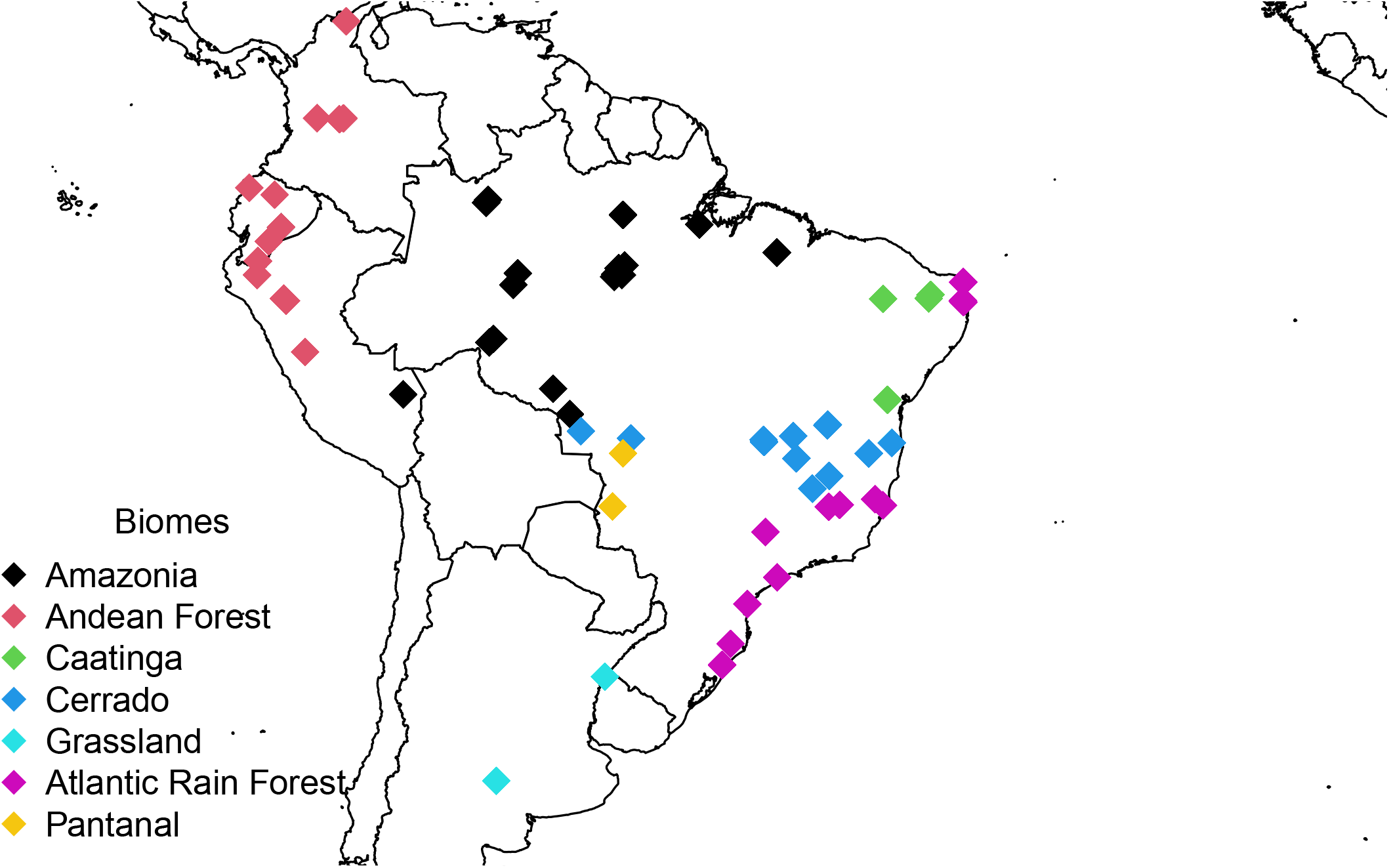
Localities where haemosporidians were sampled from birds, comprising a total of 85 localities by combining our dataset and the MalAvi database.

### Haemosporidian-Host Partner Fidelity and Network Centrality Analyses

All analyses were conducted in R version 4.0 (R Core Team, 2019). For haemosporidian-bird partner fidelity and network centrality analyses, we considered only biomes with at least 10 distinct parasite lineages, which involved 249 distinct avian host species and 40 parasite lineages from five biomes – Amazonia, Andean Forest, Cerrado, Caatinga and Atlantic Rain Forest (Supplementary Table S1). We created incidence matrices between avian host species and parasite lineages for each biome. Using the “specieslevel” function from the “bipartite” package (Dormann et al. 2008) in R, we computed normalized degree and weighted closeness and betweenness values for hosts infected by parasites in each biome. The first value represents the number of distinct realized interactions between hosts and parasites in each biome divided by the total number of distinct potential partners (i.e. parasites) in that same region. Normalized degree values can be employed as measures of partner fidelity, with hosts presenting higher values being less specific to their partners than hosts with lower values (Fortuna et al. 2020). On the other hand, weighted closeness and betweenness are measures of centrality in a network. Weighted closeness is calculated as the inverse minimum sum of the paths between a species (i.e. hosts) and all their partners (i.e. parasites) through the network, with hosts presenting higher closeness values being more central. In contrast, weighted betweenness represents the degree to which a species is positioned on the paths linking other species, i.e. the degree to which a species connects other species in an ecological network. We then combined the values obtained for birds in all biomes into one single dataset and ran a Bayesian model to compare partner fidelity and network centrality among migratory categories.

In order to run our Bayesian analyses, we employed the function “brm” from the “brms” package (Bürkner 2017). In the first model, we considered normalized degree as the response variable and avian host migratory category (resident; partial migrant and full migrant, reference level = resident) as our population-level effect and used biome as random effect. Likewise, for our second model we employed weighted closeness as the response variable, avian host migratory category (resident; partial migrant and full migrant, reference level = resident) as our population-level effect and again biome as random effect. Then, we ran a third model with weighted betweenness as our response variable, again host migratory category (resident; partial migrant and full migrant, reference level = resident) as our population-level effect and biome as random effect. We downloaded the full avian phylogeny file from the AllBirdsHackett1.tre website (https://birdtree.org/), selected only the species used for our analyses and created a matrix with phylogenetic distances between bird species. This matrix was also added to all our model as random effect to account for host phylogenetic influence on partner fidelity and network centrality. Priors were determined using the “get_prior” function. We ran the Bayesian models using 4 chains with 4000 total iterations per chain (2000 for warmup, 2000 for sampling) and employed zero-one inflated beta distributions, since normalized degree and weighted closeness and betweenness represent rate data. Further, we subsequently combined partial and full migrants into one single category and repeated our Bayesian analyses. Afterwards, we applied the “bip_ggnet” function from the “ggnet” package (<u>briatte.github.io/ggnet/</u>) to plot a bipartite net representing the relationships among haemosporidian lineages and avian hosts from different migratory categories.

### Haemosporidian Taxonomic Composition Analyses

For haemosporidian taxonomic composition analyses, we considered only localities with 10 or more individual birds sampled, at least three distinct parasite lineages per biome and at least two distinct host migratory categories, which included 2465 haemosporidian infections from 485 avian species (Supplementary Table S2). We created an incidence matrix between host migratory category and parasite lineages per biome. Later, applying the function “vegdist” (method Bray) from the “vegan” package in R (Dixon 2003), we calculated dissimilarity indices among migratory host categories. We then compared dissimilarity in parasite taxonomic composition among migratory categories using an Analyses of Variance with permutation test (PERMANOVA) for homogeneity of multivariate dispersions. For this, we employed the “permutest” function also from the “vegan” package with 999 permutations. Again, we subsequently combined partial and full migrants into one migratory category and repeated the analyses above. A non-metric multidimensional scaling plot was used to visualize the dissimilarity in parasite taxonomic composition among avian host migratory categories.

## Results

Among the 249 avian species included in the Bayesian analyses, 227 bird species were classified as resident whereas 16 and six were considered to be partially and fully migratory species, respectively. In these analyses, we assessed 81 bird species from Amazonia, 89 from Andean Forest, 73 from Cerrado, 68 from Atlantic Rain Forest and 34 from Caatinga. Our first Bayesian model revealed avian hosts display similar normalized degree (i.e. partner fidelity) among host migratory categories (Table 1) with normalized degree values around 0.10 (Fig. 2). Likewise, no difference was observed for partner fidelity when comparing resident versus non-resident (i.e. partial and full migrant hosts combined, Table 2).

**Table 1:** Parameter estimates, standard errors, and credible intervals for the Bayesian model testing the differences in partner fidelity to haemosporidian parasites among avian hosts from distinct migratory categories. (Residents only = reference category)

|  | *Estimate* | *Std. error* | *Cred. Inter (95%)* |  |
| --- | --- | --- | --- | --- |
| *Intercept* | -2.27 | 0.16 | -2.59 | -1.94 |
| *Full migratory host species* | 0.12 | 0.17 | -0.23 | 0.45 |
| *Partial migratory host species* | 0.06 | 0.11 | -0.15 | 0.27 |
| *Biomes* | 0.28 | 0.17 | 0.09 | 0.76 |
| *Avian host phylogeny* | 0.08 | 0.06 | 0.00 | 0.22 |

**Table 2:** Parameter estimates, standard errors, and credible intervals for the Bayesian model testing the differences in partner fidelity between resident and non-resident avian hosts. (Residents only = reference category)

|  | *Estimate* | *Std. error* | *Cred. Inter (95%)* |  |
| --- | --- | --- | --- | --- |
| *Intercept* | -2.28 | 0.16 | -2.60 | -1.94 |
| *Non-resident host species* | 0.08 | 0.09 | -0.11 | 0.26 |
| *Biomes* | 0.28 | 0.18 | 0.09 | 0.78 |
| *Avian host phylogeny* | 0.08 | 0.06 | 0.00 | 0.21 |

**Fig. 2:**
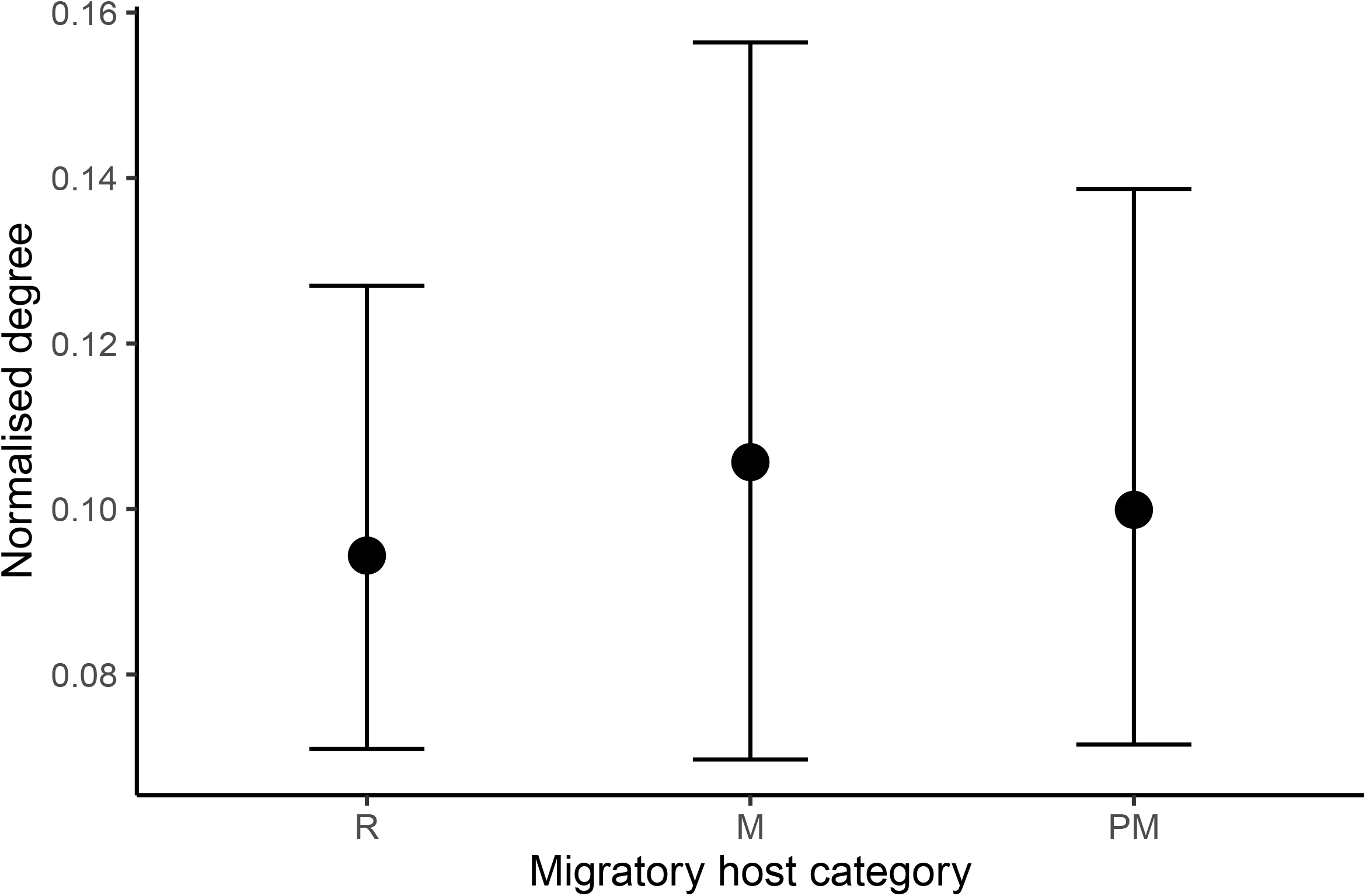
Mean (±credible intervals) normalized degree of avian hosts in bird-haemosporidian interaction networks according to the migratory category in which they are classified. R = resident, M = full migrant, PM = partial migrant.

For our next Bayesian models evaluating weighted closeness (i.e. network centrality), we observed that only partially migratory hosts present higher values of network centrality compared to residents (Table 3). On the other hand, when combining fully and partially migratory hosts into a single category, we observed that non-resident avian hosts present higher network centrality than resident species (Fig. 3, Table 4). Betweenness values were similar among host migratory categories in both models (Supplementary Tables S3, S4). Furthermore, only 51 hosts species had weighted betweenness values higher than 0, consisting of two full migratory, five partial migratory and 44 resident species. In all our Bayesian models phylogeny and biome had significant effects on partner fidelity (Tables 1-4, Supplementary Tables S3, S4). Similarly, our network plot demonstrates most hosts and parasites are found within one main component (i.e. subgroup of vertices within a graph in which there is a path possible between all vertices) and that non-resident hosts are more centrally distributed in our parasite-host network system (Fig. 4). Moreover, we can also observe that most parasites infect multiple hosts while avian hosts seem mainly infected by one or a few distinct haemosporidian lineages.

**Table 3:**
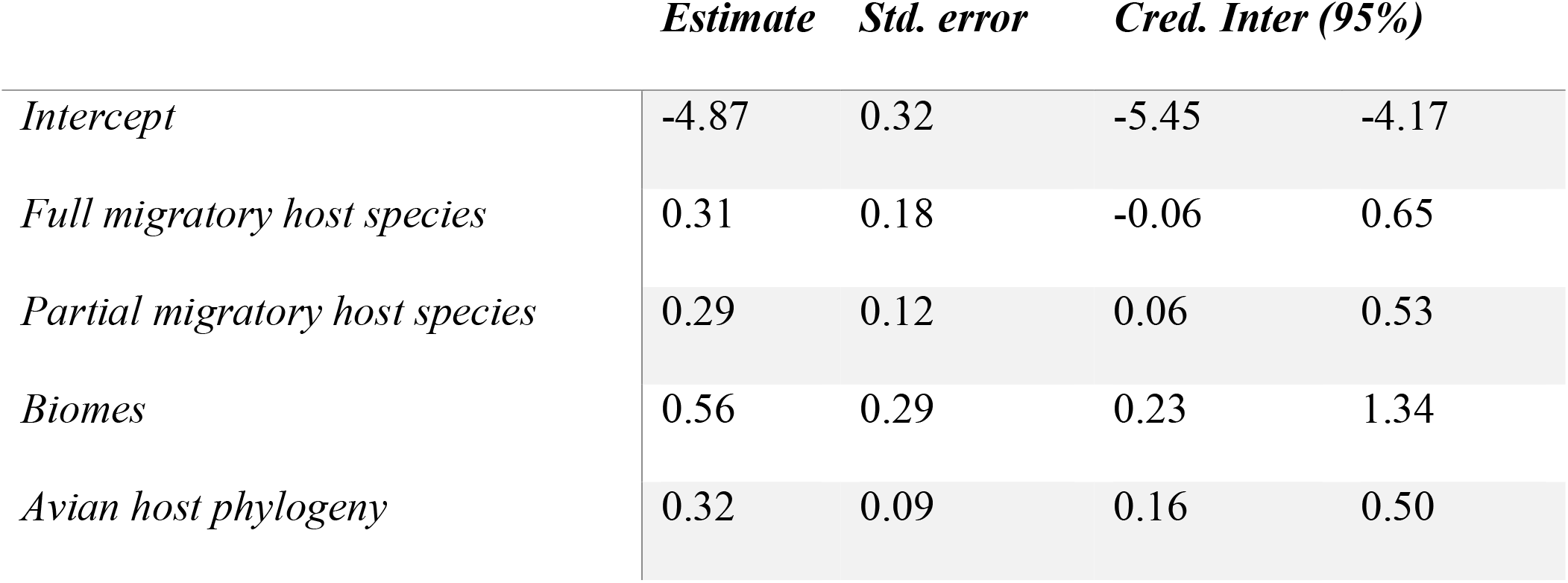
Parameter estimates, standard errors, and credible intervals for the Bayesian model testing the differences in closeness centrality to haemosporidian parasites among avian hosts from distinct migratory categories. (Residents only = reference category)

|  | *Estimate* | *Std. error* | *Cred. Inter (95%)* |  |
| --- | --- | --- | --- | --- |
| *Intercept* | -4.87 | 0.32 | -5.45 | -4.17 |
| *Full migratory host species* | 0.31 | 0.18 | -0.06 | 0.65 |
| *Partial migratory host species* | 0.29 | 0.12 | 0.06 | 0.53 |
| *Biomes* | 0.56 | 0.29 | 0.23 | 1.34 |
| *Avian host phylogeny* | 0.32 | 0.09 | 0.16 | 0.50 |

**Table 4:**
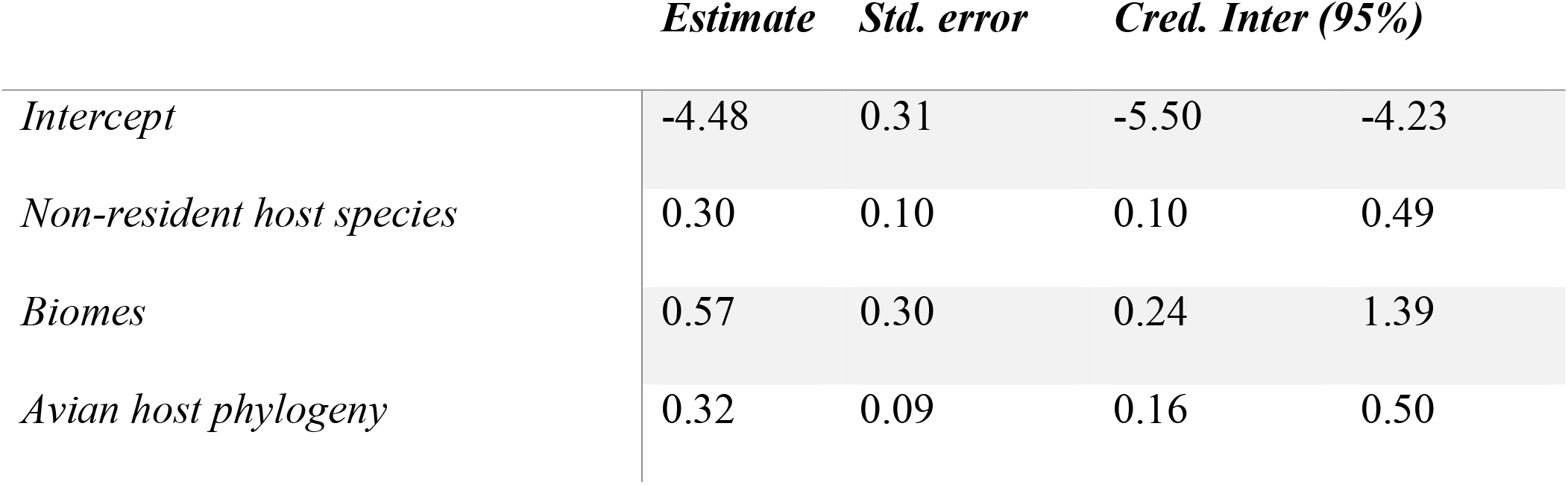
Parameter estimates, standard errors, and credible intervals for the Bayesian model testing the differences in weighted closeness of avian hosts from distinct migratory categories. (Residents only = reference category)

**Fig. 3:**
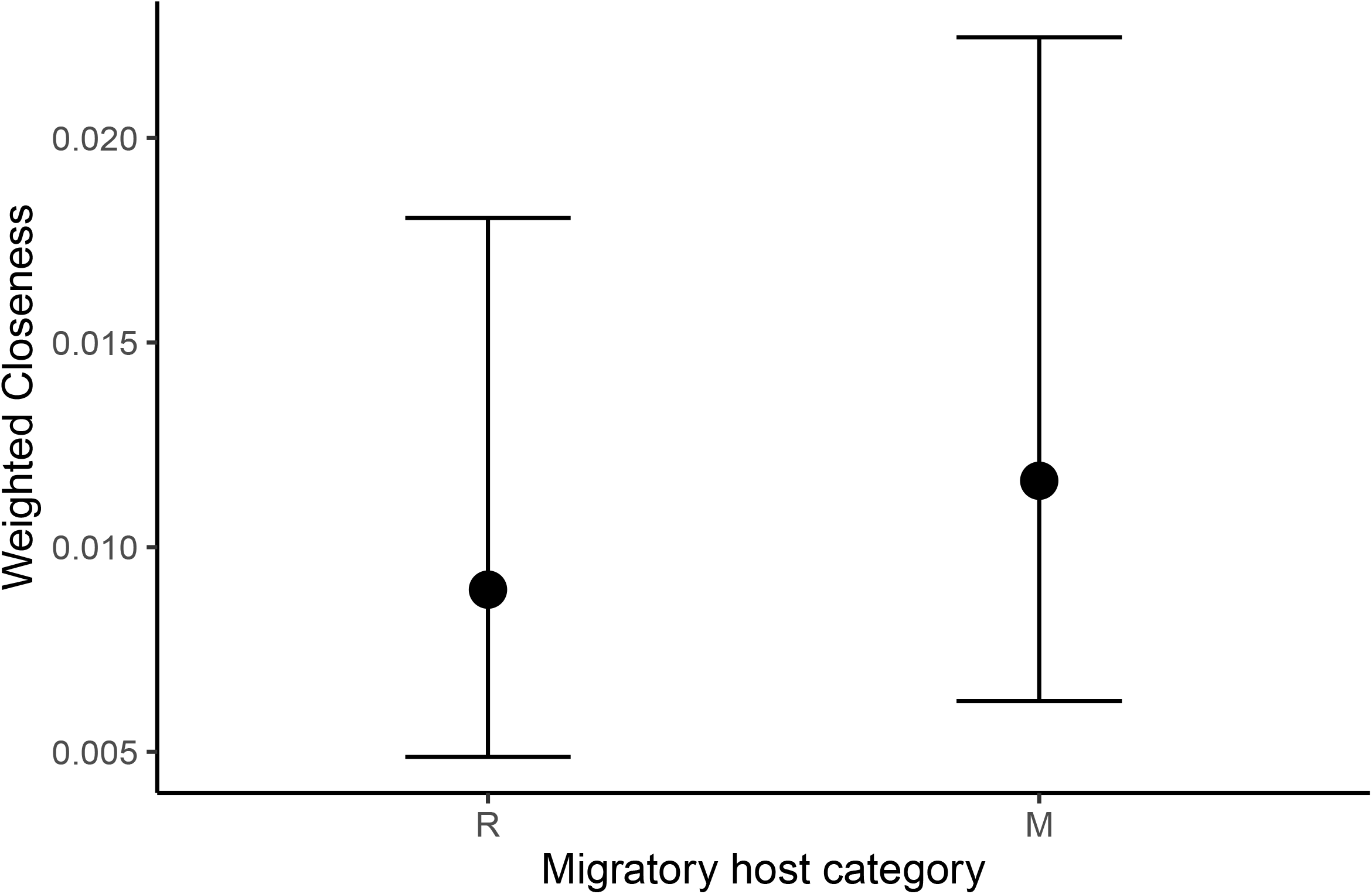
Mean (±credible intervals) weighted closeness of avian hosts in bird-haemosporidian interaction networks according to the migratory category in which they are classified. R = resident, M = full migrant and partial migrant.

**Fig. 4:**
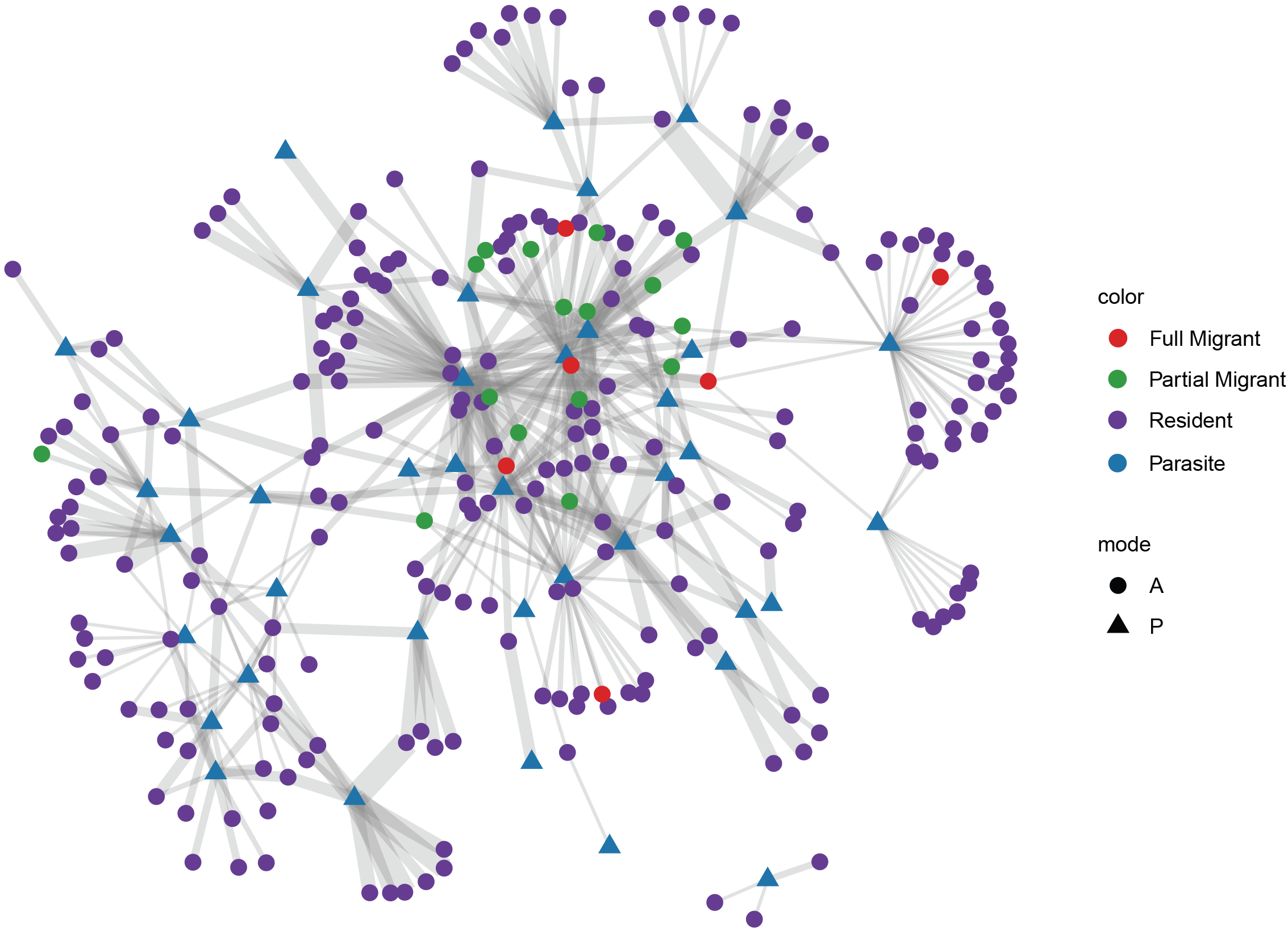
Network representing avian-haemosporidian interactions. Distinct colors represent avian hosts from distinct migratory categories or parasites. Circles represent avian hosts while triangles represent haemosporidian parasites.

Out of the 2465 haemosporidian infections included in our taxonomic composition analyses, most infections (N = 1544) represent *Plasmodium* parasites, followed by *Haemoproteus* parasites with 909, with 590 classified in the subgenus *Parahaemoproteus* and 319 in the subgenus *Haemoproteus*. Only 12 infections of *Leucocytozoon* were included in these analyses. Additionally, most parasites were recovered from Amazonia (N = 638), followed by Cerrado (N = 613) and Atlantic Rain Forest (N = 482). We observed no difference in parasite taxonomic composition among distinct migratory avian host categories when considering both resident versus partial and full migratory hosts separately (Fig. 5, Table 5) or combined (Table 6).

**Table 5:** Results of the permutational multivariate analysis of variance (PERMANOVA) testing the difference in parasite taxonomic composition among avian hosts from distinct migratory categories. (Residents only = reference category)

|  | *Degrees of Freedom* | *Sum Square* | *Mean Square* | *F value* | *P value* |
| --- | --- | --- | --- | --- | --- |
| *Groups* | 2 | 0.003365 | 0.0016823 | 0.783 | 0.46 |
| *Residuals* | 16 | 0.034376 | 0.0021485 |  |  |

**Table 6:** Results of the permutational multivariate analysis of variance (PERMANOVA) testing the difference in parasite taxonomic composition among resident and non-resident avian species. (Residents only = reference category)

|  | *Degrees of Freedom* | *Sum Square* | *Mean Square* | *F value* | *P value* |
| --- | --- | --- | --- | --- | --- |
| *Groups* | 1 | 0.000212 | 0.000212 | 0.0745 | 0.79 |
| *Residuals* | 12 | 0.034095 | 0.002841 |  |  |

**Fig. 5:**
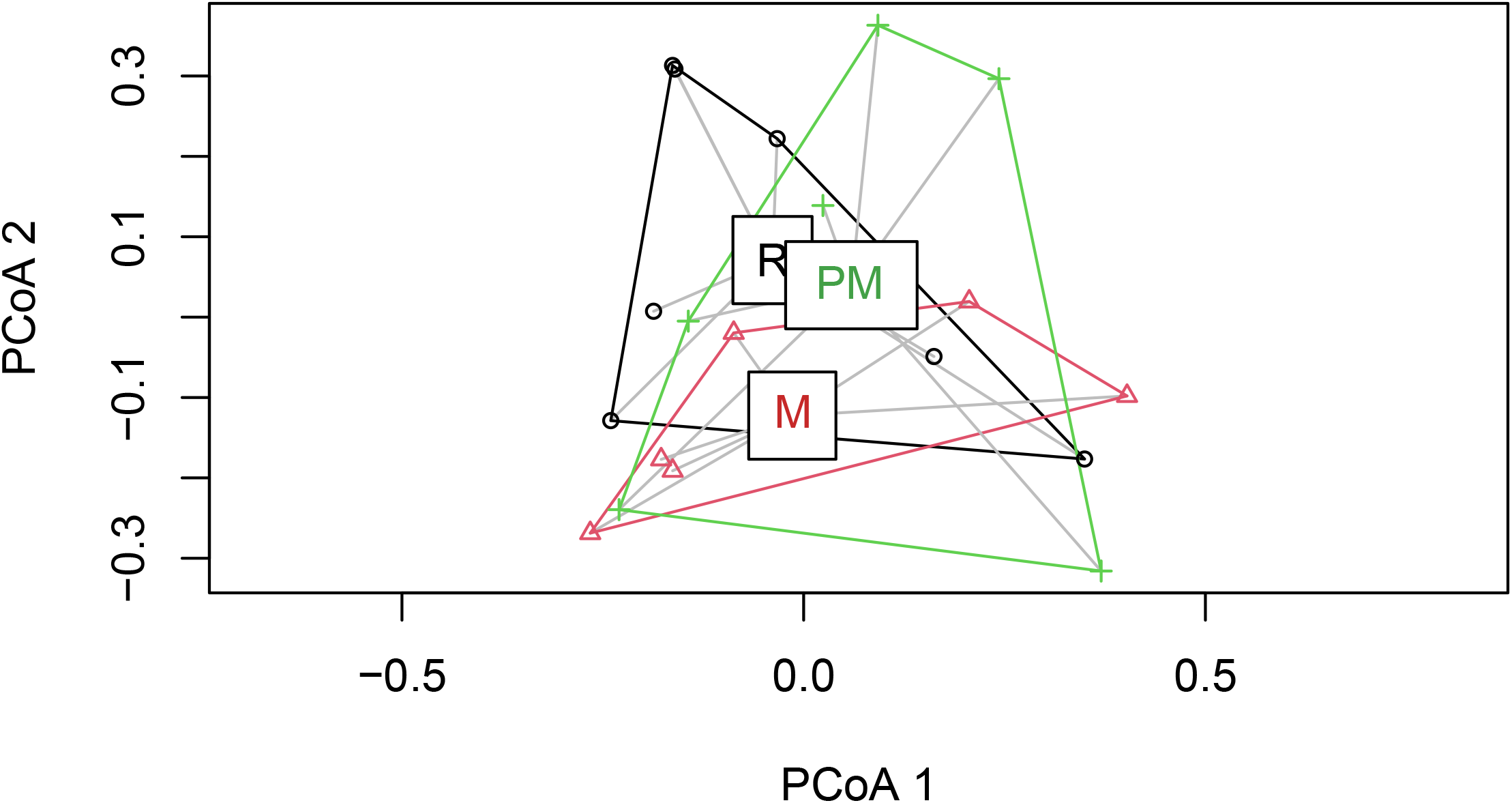
Non-metric multidimensional scaling plot illustrating the dissimilarity in parasite taxonomic composition among avian host migratory categories. R = resident, M = full migrant, PM = partial migrant.

## Discussion

Avian hosts can disperse haemosporidians across their flyways and are also able to modify local patterns of infections (de Angeli Dutra et al. 2021b), thus, migrants can play major roles into host-parasite networks. In this study, we observed that non-resident species display greater closeness centrality in host-parasites networks, which indicates they play a disproportionate role in overall network connectance (i.e. the proportion of realized interactions in a network out of the all possible interactions). However, we found no difference between resident and non-resident hosts in betweenness centrality and that most species are not network connectors (betweenness centrality = 0). This result suggests that, despite the fact migrants drive overall network connectance, these hosts do not necessarily act as key connectors between species within the network. Additionally, we also observed that resident and non-resident hosts show similar partner fidelity and parasite taxonomic composition, demonstrating that similar parasites infect resident and non-resident hosts and that there is no difference in pairwise parasite specificity among migratory and non-migratory species.

By connecting more species within the network, migratory hosts can act as keystone species (i.e. species with disproportionate importance in keeping the structure and ecological services and functions within a community; *sensu* Paine 1969) since they interact with more distinct parasite lineages and are more closely associated with further hosts. Therefore, the presence of migrants in a community could impact local parasite-host dynamics. Indeed, previous research has associated the presence of migratory birds with variation in tick prevalence and haemosporidian prevalence and richness within the local community in South America (de Angeli Dutra et al. 2021b; Fecchio et al. 2021). In contrast, despite the fact that only partially migratory hosts presented higher closeness centrality when evaluated separately, de Angeli Dutra et al. (2021a) observed that only fully migratory birds harbor higher prevalence and richness of haemosporidian parasites. Nevertheless, no difference was observed here with respect to betweenness centrality, suggesting resident and non-resident hosts play similar roles in connecting parasites and other hosts. Thus, since migrants show higher closeness centrality and are involved in disproportionately more interactions within the network, they are influential in shaping parasite transmission within the community.

We also demonstrated that migration does not impact partner fidelity for haemosporidian parasites and their avian hosts. Hence, it is possible the predictability of migration patterns allows parasites to co-adapt to these hosts as successfully as they do for resident species. Furthermore, the trade-off between adapting to multiple environments and vectors may be compensated by the opportunities to colonize new habitats and host species provided by host migration. Concomitantly, haemosporidian parasites tend to infect wide subsets of phylogenetically related avian hosts (Pinheiro et al. 2016; Huang et al. 2018). Thus, parasite host specificity patterns may remain similar within subsets of hosts which include resident and non-resident species, leading to similar parasite fidelity and taxonomic composition among distinct migratory categories. Indeed, we observed a host phylogenetic effect in all our Bayesian models, indicating that host phylogeny may be associated with multiple factors shaping host-parasite networks. Furthermore, similarity in environmental conditions also seems to affect network structure for parasites and their hosts as biome category (included as a random factor) also influenced partner fidelity and centrality in all our models. Likewise, previous research suggests that climate variation is an important driver of haemosporidian parasite specificity in South America (Fecchio et al. 2019b). Therefore, host phylogeny and environment may be better predictors of parasite fidelity and taxonomic compositions than host migratory behavior.

Antagonistic interactions are generally characterized by lower partner fidelity patterns and, therefore, more malleability than mutualistic interactions (Fortuna et al. 2020). Therefore, parasites may be associated with looser evolutionary pressures for specialization favoring colonization of new habitats and spillover events. Indeed, a recent spillover of *Plasmodium juxtanucleare* from domestic and exotic hosts (chickens) to wild passerine birds has been reported in Brazil (Ferreira-Junior et al. 2018), demonstrating haemosporidian parasites can adapt to new hosts when placed in alien habitats. Moreover, Krasnov et al. (2012) argued that parasites can infect unrelated hosts when phylogenetically close hosts are exploited by too many pathogens. These findings suggest that parasites are malleable enough to exploit unfamiliar hosts in response to adverse resource conditions. This plasticity could lead to looser interaction patterns in avian-haemosporidian networks and similar dynamics for resident and non-resident birds. Nevertheless, host-parasite networks tend to be compartmentalized into modules (Bascompte 2010; Krasnov et al. 2012), which may reflect an ongoing arms race between parasites and their hosts (Bascompte 2010) and consequential convergence of traits among distinct parasites (Krasnov et al. 2012).

In summary, we show migratory hosts may be keystone species within host-parasite networks and their presence could putatively shape bird-haemosporidian interactions by, for example, impacting local prevalence and richness of parasites (de Angeli Dutra et al. 2021b). Additionally, most birds are not important connectors in this network, with resident and non-resident hosts playing similar parts in connecting hosts and parasites. However, it is important to note that, despite the fact most avian hosts are not network connectors, most species belong to a single network component. Moreover, no difference in partner fidelity or parasite taxonomic composition was detected in this study between migrant and non-migrant birds, indicating parasite specificity may be associated with other traits of avian and vector hosts. Further, biome and phylogeny seem to play important roles in determining network characteristics of hosts in avian-haemosporidian networks, an effect already demonstrated in systems involving trophically transmitted parasites (Poulin et al. 2013). We conclude that migrants may play fundamental roles in shaping host-parasite interactions, and encourage further research into other potential implications of host migration for disease dynamics.

## Acknowledgments

We thank the MalAvi curators for maintaining the database and for making all data available, as well as all researchers who deposited their data into this public repository. We are also grateful to Lucas Marques for graphical support.

## Availability of data and material

A part of the data that support the findings of this study is openly available at https://onlinelibrary.wiley.com/doi/10.1111/mec.15094 and http://130.235.244.92/Malavi/ (Bensch et al. 2009). The other portion of the data that support our findings can be shared by Prof. Érika Martins Braga under reasonable request.

## Authors’ contribution

Daniela Dutra and Robert Poulin conceived the idea and designed the study. Daniela Dutra performed the data analyses. Daniela Dutra, Érika Braga and Alan Fecchio collected the data. Daniela Dutra wrote the manuscript with input from all other authors. All authors contributed critically to the drafts and gave final approval for publication.

