## Supplementary Table 3-4 for "Haemosporidian taxonomic composition, network centrality and partner fidelity between resident and migratory avian hosts"

**Electronic Supplementary Material:**

Supplementary Table 3: Parameter estimates, standard errors, and credible intervals for the Bayesian model testing the differences in weighted betweenness among avian hosts from distinct migratory categories. (Residents only = reference category)

|  | **Estimate** | **Std. error** | **Cred. Inter (95%)** | |
| --- | --- | --- | --- | --- |
| Intercept | -2.43 | 0.36 | -3.18 | -1.70 |
| Full migratory host species | 0.79 | 0.64 | -0.58 | 1.96 |
| Partial migratory host species | 0.13 | 0.48 | -0.85 | 1.01 |
| Biomes | 0.29 | 0.27 | 0.01 | 0.98 |
| Avian host phylogeny | 0.53 | 0.28 | 0.05 | 1.13 |

Supplementary Table 4: Parameter estimates, standard errors, and credible intervals for the Bayesian model testing the differences in weighted betweenness between resident and non-resident avian hosts. (Residents only = reference category)

|  | **Estimate** | **Std. error** | **Cred. Inter (95%)** | |
| --- | --- | --- | --- | --- |
| Intercept | -2.42 | 0.36 | -3.15 | -1.69 |
| Non-resident host species | 0.37 | 0.40 | -0.46 | 1.14 |
| Biomes | 0.30 | 0.28 | 0.01 | 1.07 |
| Avian host phylogeny | 0.53 | 0.28 | 0.04 | 1.16 |
